# Detecting single motor-unit activity in magnetomyography

**DOI:** 10.1101/2024.10.15.618402

**Authors:** Nima Noury, Justus Marquetand, Stefan Hartwig, Thomas Middelmann, Philip Broser, Markus Siegel

## Abstract

Studying the discharge patterns of motor units (MUs) is key to understanding the mechanisms underlying human motor behavior. Intramuscular electromyography (iEMG) allows direct study of MU activity, but is invasive. Surface electromyography (sEMG) offers a non-invasive alternative, but with lower spatial resolution. Recent advances in optically pumped magnetometers (OPMs) have sparked interest in the magnetic counterpart of EMG, magnetomyography (MMG), as an additional non-contact modality to study the neuromuscular system. However, it remains unclear whether MMG signals recorded with superconducting quantum interference devices (SQUIDs) or OPMs can be used to directly detect individual MUs. We addressed this question in a proof-of-principle study in which we recorded MMG signals from the abductor digiti minimi (ADM) muscle using SQUIDs and OPMs. Critically, we simultaneously recorded iEMG from the same muscle to validate the non-invasive measurements. First, we found that invasively recorded MUs can be detected in simultaneously recorded SQUID and OPM MMG signals. Second, we found that invasively validated MUs can be extracted directly from SQUID and OPM MMG. This provides converging evidence that individual MU activity is accessible using non-contact MMG. Our findings highlight the potential of MMG as a non-contact modality to measure and study muscle activity in health and disease.

## Introduction

Studying the discharge patterns of motor units is key to understanding the mechanisms underlying human movement^1–3^. Each motor unit (MU) is comprised of a motor neuron and all muscle fibers that are innervated by that motor neuron^4^. The innervated muscle fibers amplify the comparatively small action potentials of motor neurons into much bigger motor unit action potentials (MUAPs). Thus, each firing of a motor neuron results in a comparatively large motor unit potential that can be recorded through electromyography (EMG)^3^. In turn, studying motor unit activity can be considered an indirect way to study motor neuron activity at the output stage of the spinal cord.

The most direct way to study MU activity is by intramuscular EMG (iEMG), i.e., by invasively measuring bioelectrical activity through needles or fine-wire electrodes inserted directly into the muscle fibers^1,3^. Despite the high spatiotemporal resolution iEMG also comes with disadvantages. One limitation is that iEMG reflects local activity and, therefore, it is difficult to study the orchestrated activity of several different motor unit populations. Another limitation of iEMG is its invasive nature, which is inconvenient and sometimes even impossible to withstand for subjects and patients^5^. In fact, in some countries, medical and technical constraints restrict the use of iEMG^6^.

Electrical currents that result from MU activity lead to voltage differences on the skin around the muscle. Surface EMG (sEMG) captures these voltage differences on the skin. but has a much lower spatial resolution as compared to iEMG^7,8^. The signal of a single sEMG electrode reflects the summed activity of a population of MUs up to centimeters away from the sEMG electrode^9^. Advances in signal processing have made it possible to decompose recordings of sEMG electrode grids into underlying MU activities^6,7,10–17^. This approach has been useful not only for studying motor units, but also for human-machine interfaces^6,18–21^.

An entirely non-invasive, i.e. contactless alternative to EMG, is to measure the magnetic counterpart of electrical MU activity – magnetomyography (MMG). MMG was introduced in the 1970s^22^. MMG recorded with superconducting quantum interference devices (SQUID) reflects the activity of MUs detected in simultaneous sEMG recordings^23^. However, MMG has not been widely utilized for studying human muscle activities^24,25^. This is primarily due to the high cost and bulkiness of SQUID technology, which requires cryogenic cooling and fixed sensor array geometries. These technical constraints have been overcome by recent advances in quantum sensing through the development of optically pumped magnetometers (OPMs). OPM can be flexibly placed around all body parts and muscles due to their miniaturized design and lack of cryogenic cooling^26^. Consequently, utilizing MMG for the study of human muscles has recently gained attention^5,9,27–34^.

Biophysical simulations suggest that some MMG components exhibit a better spatial selectivity and are less influenced by subcutaneous fat than sEMG^9^. Along the same line, it was demonstrated in-silico that high-density MMG (HD-MMG) could be superior to high-density sEMG for motor unit decomposition (HD-sEMG)^30^. However, these simulations have been conducted under ideal scenarios, and, as of now, no *in vivo* experiment has addressed the question if motor units can be directly decomposed from MMG recordings. Therefore, it remains unclear if MU decomposition from MMG recordings is possible with current technologies. This is especially important for MMG signals recorded with OPM, because with a maximal bandwidth of 2 kHz and a critical bandwidth-sensitivity tradeoff^35^ currently available OPM-technologies are not comparable to sEMG (10 kHz typical bandwidth).

To address these questions in a proof-of-principle study, we recorded multichannel MMG signals with SQUID and OPM technologies. We tested if single MUs can be directly identified from these MMG recordings. Critically, we validated MU identification through simultaneous iEMG recordings. Our results show that single MU activity can be identified from both SQUID and OPM MMG.

## Materials and Methods

### Participants and experimental protocol

Two healthy subjects participated in this study and provided written informed consent. The study was carried out in accordance with the Declaration of Helsinki and was approved by the ethics committee of the University of Tübingen.

### Data acquisition

We simultaneously recorded intramuscular EMG (iEMG) and MMG. For each subject, we used either SQUID sensors or OPMs to record MMG. SQUID recordings were done at 2.3438 kHz sampling rate at the MEG center Tübingen, Germany, using a 275-channel MEG system (Omega 2000, CTF Systems). This MEG system also provided an integrated EEG recording interface, which we used to record simultaneous iEMG. OPM recordings were performed at 8 kHz sampling rate using five zero-field OPMs (triaxial QZFM-gen-3; QuSpin Inc., Louisville, CO, USA) at Berlin magnetically shielded room-2 (BMSR-2), Physikalisch-Technische Bundesanstalt (PTB), Berlin, Germany^26^. During OPM recordings, simultaneous iEMG was recorded using a NeurOne EEG system (Mega Electronics Ltd) at 20 kHz sampling rate. For iEMG, we used custom-made non-magnetic concentric bipolar needle electrodes with a recording area of about 0.07mm^2^ comparable to clinical neurophysiological standard (depth roughly equals 1 cm). Electrodes consisted of a platin-shaft and a platin-iridium wire inside the shaft. All iEMG-recordings were performed by an experienced board-certified clinical neurophysiologist (JM). In the OPM experiment, we measured ADM abduction force and the distance between the hand and OPM lateral to the ADM using custom devices. Force and distance signals were recorded using the same NeuroOne recording setup as the iEMG signals.

In each subject, we recorded the activity of the abductor digiti minimi muscle (ADM) during 60 s intervals of rest, light abduction, and maximum abduction. For light and maximum abduction conditions, subjects constantly abducted their pinky finger, but their hands and fingers were not fixed. For the SQUID recordings, we positioned the MEG dewar horizontally and placed the subject”s hand inside the MEG helmet (Figure 1). OPM were positioned on a half-circle at the hand palm”s transverse plane and distanced 45 degrees from each other (Figure 1).

**Figure 1.**
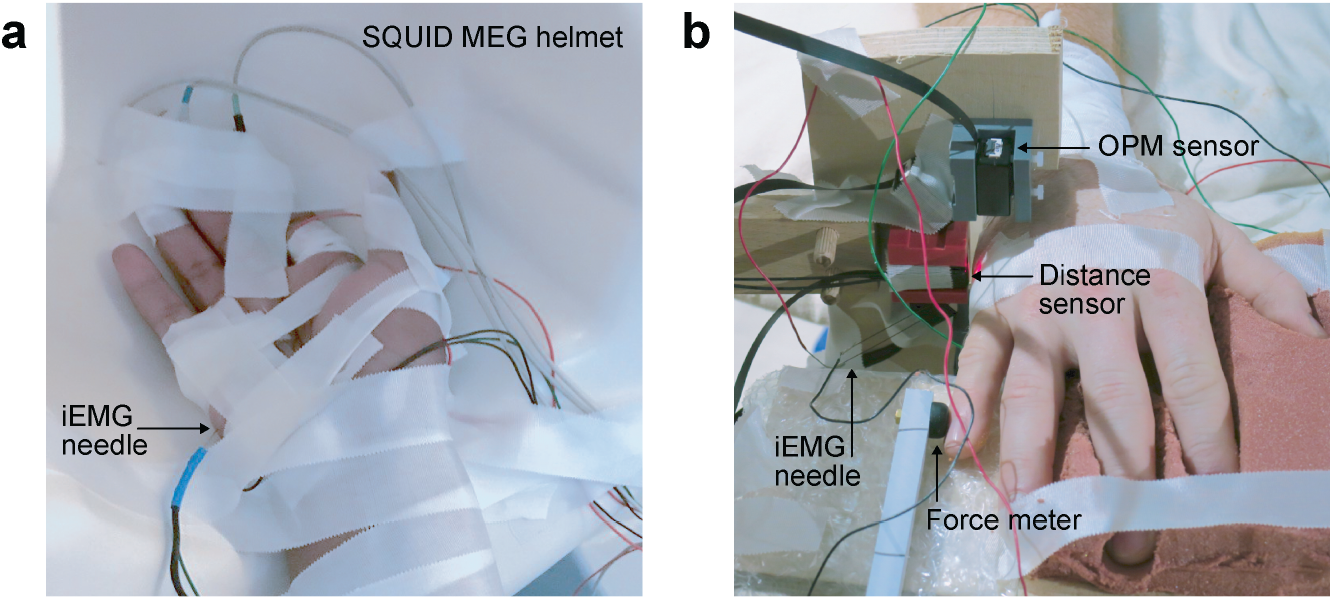
Experimental setups. **a**, SQUID setup for experiment 1. The SQUID-MEG dewar (CTF Systems) was positioned horizontally and the subject”s hand was placed on the inner rear side of the MEG helmet. **b**, OPM setup for experiment 2. 5 triaxial OPM sensors (QuSpin Inc.) were positioned on a half-circle around the ADM muscle distanced 45 deg from each other. During the OPM experiment, abduction force and hand distance to one of the OPM sensors were measured continuously.

### Preprocessing

Because of the variable sampling rates and information content of different modalities, we adapted the preprocessing for each recording. For the SQUID experiment, we high-pass filtered the iEMG at 10 Hz and band-pass filtered the SQUID signals from 10 to 900 Hz. For the OPM experiment, we band-pass filtered the iEMG and OPM signals from 20 to 2000 Hz and from 20 to 800 Hz, respectively. All filters were 4^th^ order Butterworth filters. We notch filtered line noise up to the 10^th^ harmonics using 2^nd^ order Butterworth band-stop filters with 1 Hz bandwidth. Even after filtering, the OPM signals showed a technical artifact around 923 Hz^34^, which was attenuated using 2^nd^ order Butterworth band-stop filters from 910 to 936 Hz. For the OPM experiment, iEMG and MMG signals were down-sampled by a factor of 3 resulting in 20/3 kHz and 8/3 kHz, respectively. With exception of the MUAPs and MUAFs (motor unit action fields) shown in Figures 5 and 6, all filters were applied forward and backward (i.e., zero phase filtering). For these exceptional cases, we observed that zero phase filtering together with the low bandwidth of OPM signals in comparison to the very high bandwidth of iEMG led to the misleading impression that some OPM MUAFs started earlier than iEMG MUAPs. Thus, for generating the curves depicted in Figures 5 and 6, we expanded the OPM band-pass filter to 10 to 900 Hz and used only forward filtering. A control analysis of the MU comparison (see below) on the OPM signals as filtered for display (forward only filter) yielded the same results as for the original preprocessing (forward-reverse filter). For SQUID MUAFs, we did not observe any significant change in curves due to zero phase filtering compared to forward filtering.

From the 272 recorded SQUID channels, we selected 40 channels that clearly reflected muscle activity. For the iEMG signal and each SQUID channel, we concatenated the signals recorded during rest, light abduction, and maximum abduction and calculated the signal envelope using the Hilbert transform. Next, we low-pass filtered the envelopes with a 2^nd^ order Butterworth filter at 0.25 Hz (green lines in Figure 2a) and correlated each SQUID signal envelope with the envelope of the iEMG signal. We then selected the 40 SQUID channels with highest correlation for further analysis.

**Figure 2.**
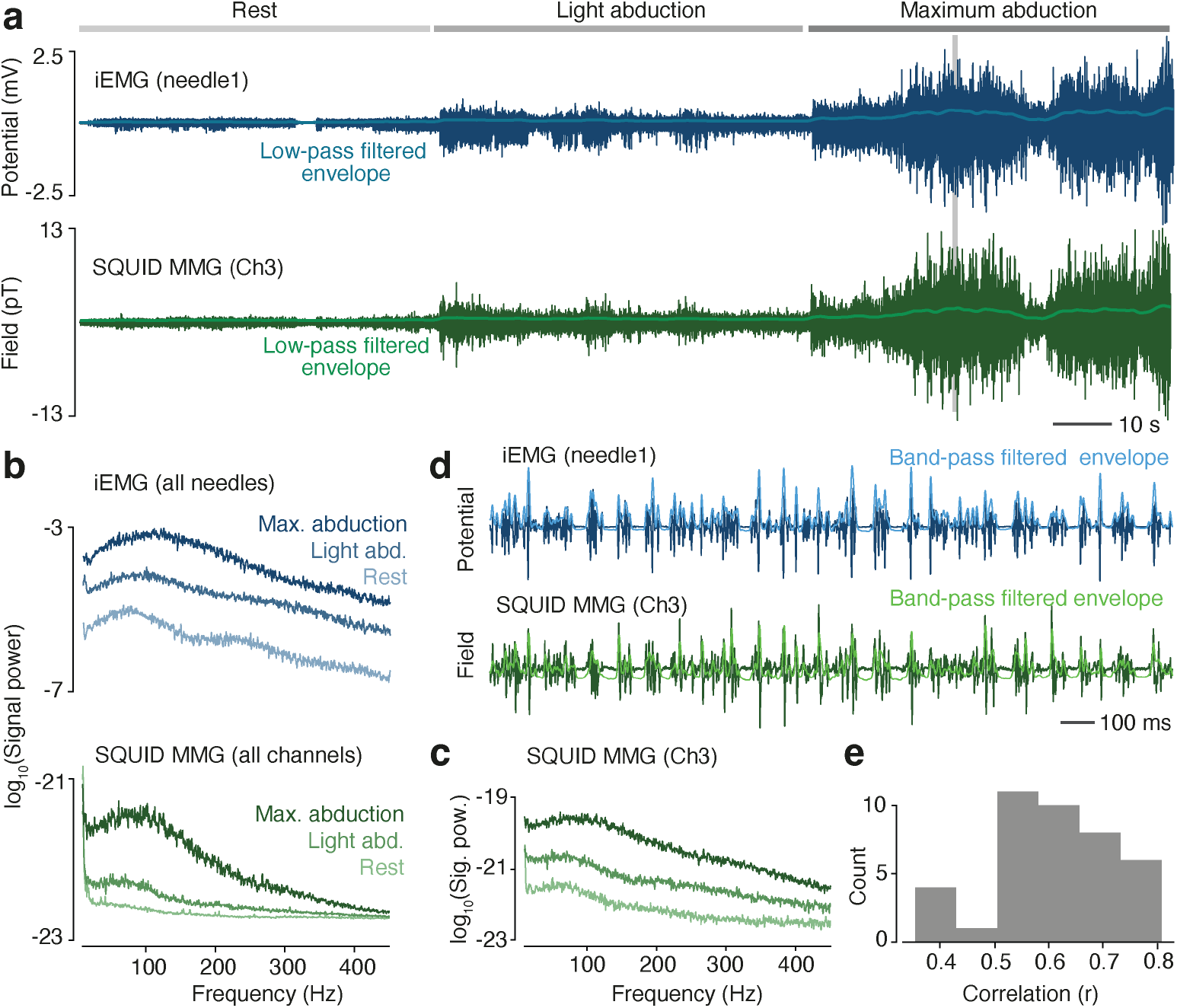
Experiment 1: SQUID MMG and iEMG. **a**, iEMG (top) and simultaneously recorded SQUID MMG. Bright lines show the low-pass filtered signal envelopes (0.25 Hz cut-off). The gray area indicates the 2 s interval shown in (d) **b**, Power spectra of iEMG (top) and SQUID MMG (bottom) averaged across sensors. **c**, Power spectra of the MMG channel shown in (a) and (d). **d**, 2 s of iEMG and MMG. Bright lines show the band-pass filtered signal envelopes (1 to 100 Hz). **e**, Histogram of the correlation between the high-pass filtered envelopes of iEMG and SQUID MMG across the 40 SQUID channels that were selected, based on low-pass filtered envelopes.

### Decomposition

We applied convolutive ICA to decompose iEMG and MMG recordings into MUs ^14^ using the open-source toolbox emgdecomp^21^. We used 15 ms and 10 ms extension factors for MMG and iEMG decomposition, respectively. Silhoutte Score (SIL) thresholds were 0.85 for iEMG and SQUID signals and 0.9 for OPM signals. For estimating the percentage of overlapping firings we considered a maximum lag of 25 ms between spike trains and a jitter of +/-0.5 ms for each spike^14^.

We tested the significance of kernels (i.e. MUAPs and MUAFs in Figures 3 and 5) using a permutation statistic. For each component and channel, we calculated the 50 ms long average kernel across firings and compared its temporal standard deviation against a distribution of 1000 standard deviations generated under the null-hypothesis. Null-hypothesis standard deviations were generated by randomly jittering each firing moment (uniform distribution between -1 and 1 second) before calculation the average kernel and its temporal standard deviation.

**Figure 3.**
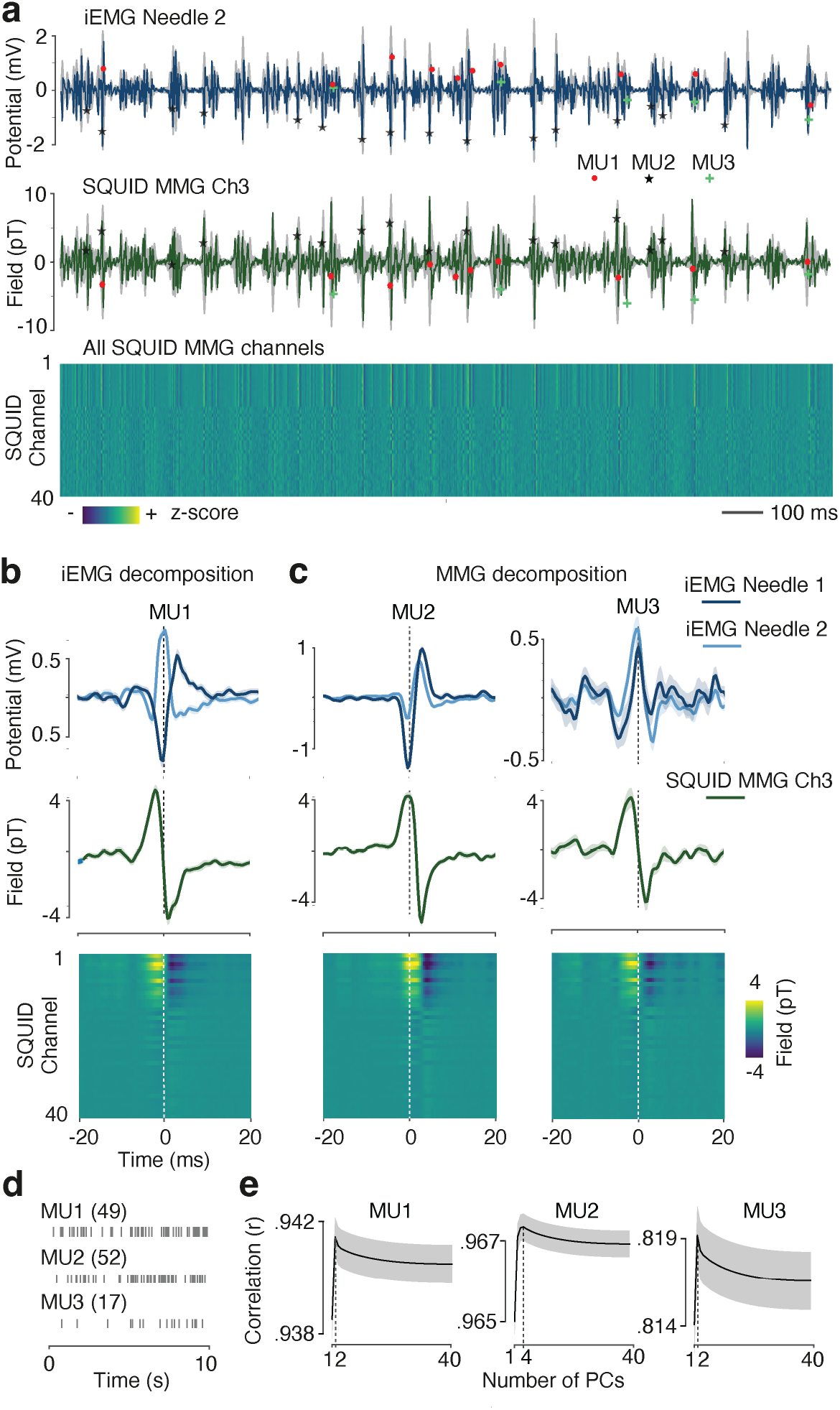
MU detection in experiment 1. **a**, 2 s of iEMG (top) and SQUID MMG (middle and bottom). Markers indicate the firing moments of the 3 discovered MUs. **b**, MUAP (top) and MUAF (middle and bottom) of the MU decomposed from iEMG. **c**, MUAPs (top) and MUAFs (middle and bottom) of the 2 MUs decomposed from SQUID MMG. **d**, Firings of the 3 decomposed MUs in the first 10 s of the recording. The number of firings is in brackets. **e**, Dimensionality estimation of the 3 MUAFs by cross-validated PCA. All shaded regions depict SEM.

### Comparing MU activation patterns

We compared activation patterns of MUs using LDA classifiers. For each MU, we generated a population of waveforms by selecting the 50 ms intervals around all firing moments. For example, for a MU with 100 firings, in an MMG recording with 40 channels and sampling rate of 1 kHz, this resulted in a 100 × 40 × 50 matrix, corresponding to one row of panels in Figures 4c, 4d, 6d, and 6e.

**Figure 4.**
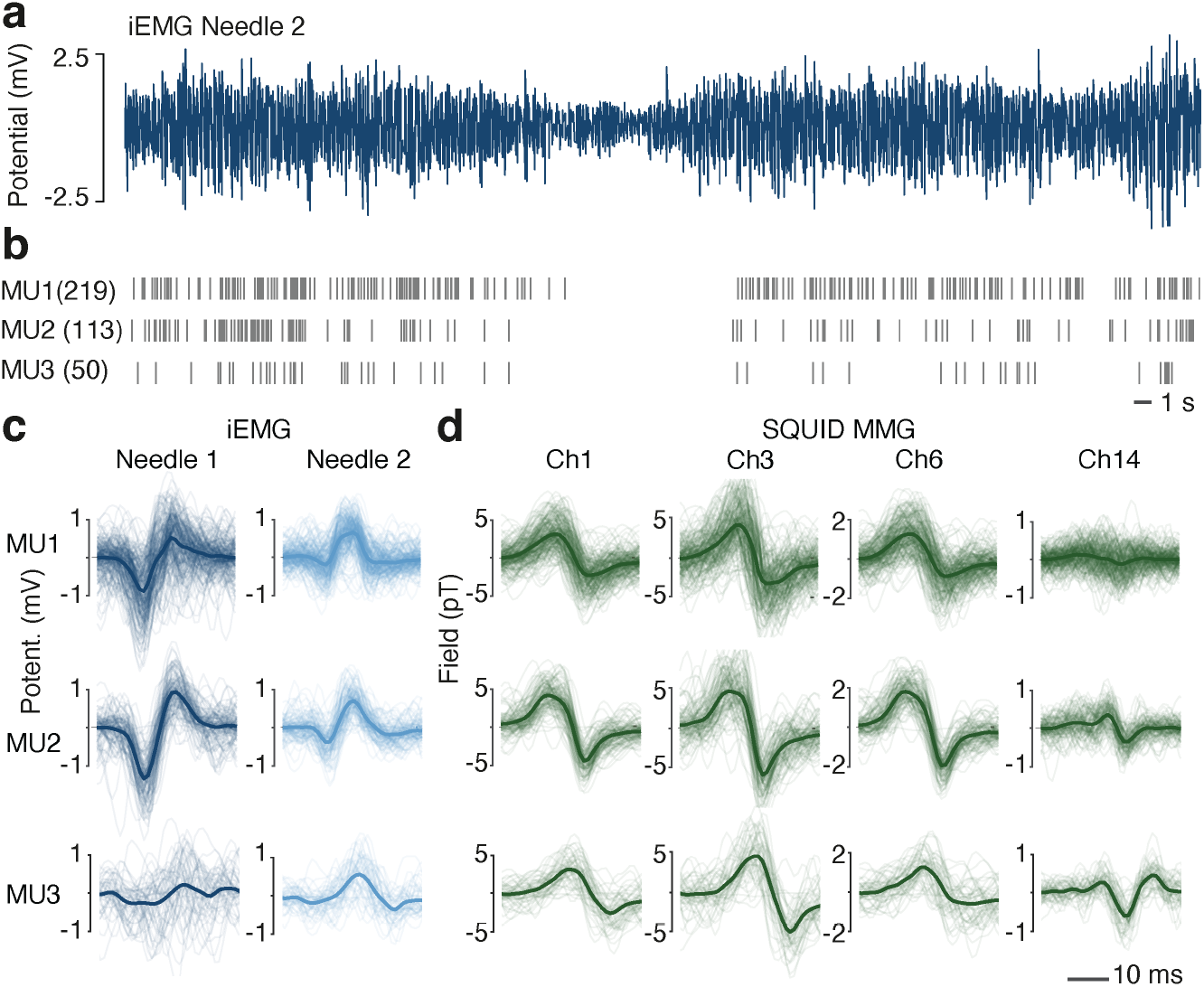
Dissociating MUs in experiment 1 **a**, top: iEEG during the entire 60 s of maximum abduction. Bottom: firing moments of the 3 identified MUs. MUs were decomposed from the first 10 s. Discharge patterns of MUs reflect the amplitude of the IEEG. **b**, single discharge (thin lines) and average (thick lines) iEEG MUAPs of all identified MUs. iEEG was band-pass filtered from 20 to 1000 Hz (zero-phase filtering). **c**, single discharge (thin lines) and average (thick lines) SQUID MUAFs (4 representative channels) of all identified MUs. MMG signals are band-pass filtered from 10 to 900 Hz (zero-phase filtering).

To compare two MUs, we first temporally aligned these two MUs. We first calculated the average signal of each MU across firings and sensors and determined the latency of the global peak and trough. Then, we chose as reference MU the wider MU, i.e. the MU with a larger delay between peak and trough. We cropped all waveforms of the reference MU to a 10 ms long interval around the center between peak and trough. Then, we aligned the second MU to the reference MU. We slid a 10 ms long window over the second MU, and, for each window and each sensor, we calculated the average signal across all firings. We then determined the 10 ms interval with the highest correlation between the cropped reference MU and the second MU. For calculating this correlation, we concatenated average signals of all channels of each MU and calculated the Pearson correlation of these two vectors.

We then compared the aligned waveforms using an LDA classifier. To account for different number of firings across MUs, we balanced the population by randomly choosing the same number of firings of the MU with more firing as the number of firings of the MU with less firings. Then, we trained a 10-fold cross-validated LDA on these two populations and calculated its F1-score. We employed a permutation statistic to estimate statistical significance. We compared the measured F1-score to a distribution of 1,000 F1-scores generated under the null-hypothesis to obtain the one-tailed p-value. To generate the null distribution, we randomly shuffled the firing labels (i.e., randomly assigned each firing to one of the MUs), calculated the F1-score as described above, and repeated this process 1,000 times.

We performed a control analysis to test if our approach yielded spurious results. We randomly divided all 184 firings of MU2 of the OPM experiment (Figure 6) into two groups and shifted the waveforms of one of these groups by 10 ms. We then applied the above analysis pipeline to compare these two pseudo-components. As expected, this resulted in no significant difference (F1-score = 0.51, p = 0.44).

### PCA analysis

We used PCA to estimate the dimensionality of MUAFs (panels b, c, and e in Figures 3 and 5). We randomly divided firings of each MU into two groups and averaged across firings to obtain two kernels. We applied PCA across n channels of the first kernel, and generated n dimension-reduced kernels, in which the m-th version contained only PCs 1 to m. We concatenated signals of all channels and correlated the second kernel with each of the n dimension-reduced versions of the first kernel. We repeated this procedure 1000 times. Then, for each m, we calculated the average of the 1000 correlations and found the m with the highest average correlation as the dimensionality of the MMG kernel (Figure 3e).

**Figure 5.**
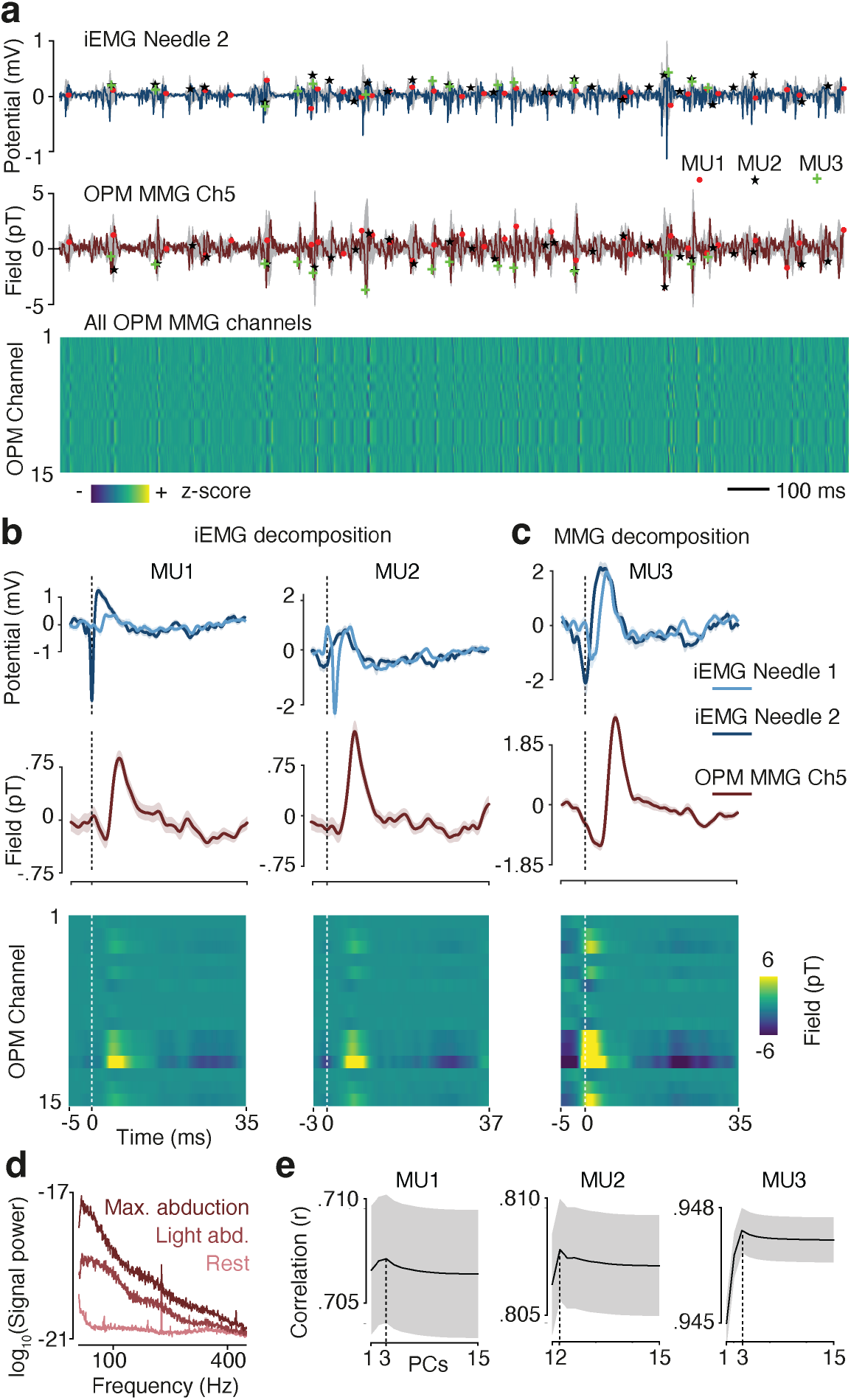
MU detection in experiment 2. **a**, 2 s of iEMG (top) and OPM MMG (middle and bottom). Markers indicate the firing moments of the 3 discovered MUs. **b**, MUAP (top) and MUAF (middle and bottom) of the 2 MUs decomposed from iEMG. **c**, MUAPs (top) and MUAFs (middle and bottom) of the MU decomposed from OPM MMG. **d**, Power spectra of OPM MMG averaged across sensors. **e**, Dimensionality estimation of the 3 MUAFs by cross-validated PCA. All shaded regions depict SEM.

### Analysis software

All data analyses were performed in Python using custom scripts and the open source toolbox emgdecomp^21^.

### Data and materials availability

The data that support the findings of this study are available from the corresponding authors upon reasonable request.

## Results

### SQUID MMG

In a first experiment, we simultaneously recorded SQUID MMG and iEMG, to test if single MUs can be decomposed from SQUID MMG. (Figure 2). We recorded the activity of the abductor digiti minimi muscle (ADM) of the hand of one subject during 60 s intervals of rest, light and maximum finger abduction. MMG recordings were performed with a 272-channel whole-head MEG system placing the subject”s hand inside the MEG helmet (Figure 1a). As expected, with increasing voluntary muscle contraction, the iEMG showed a robust increase of signal variance in the time domain (Figure 2a) corresponding to a broadband power increase in the frequency domain (Figure 2b). Similar to iEMG, also the variance of individual SQUID channels (Figure 2a) and the average power across all SQUID channels (Figure 2b) showed a broadband increase with increasing muscle contraction. Because of the sensor geometry of the MEG-helmet not all SQUID channels were placed near the hand. Thus, for further analyses, we identified channels that well reflected ADM muscle activities. To this end, we correlated the low-pass filtered envelopes (0.25 Hz cut-off) of iEMG and MMG signals (bright lines in Figure 2a) and selected the 40 SQUID channels with highest correlation to the iEMG (average Pearson”s correlation coefficient of selected channels = 0.93 ± 0.06, correlation of channel shown in Figure 2 = 0.997). When we zoomed into the time domain, we also observed a high degree of similarity between spike-like events in band-pass filtered iEMG and SQUID MMG signals (Figure 2d; 10 to 500 Hz band-pass). Indeed, also the band-pass filtered envelopes (bright lines in Figure 2d; 1 to 100 Hz) of band-pass filtered iEMG and of the 40 selected MMG channels were highly correlated (Figure 2e, correlation of channel shown in Figure 2d = 0.72; p < 1e-6). In sum, we observed strong similarities between SQUID and iEMG signals.

Next, we investigated whether the activity of single MUs could be measured with SQUIDs. To address this, we first identified a single MU in the iEMG and then tested if the SQUID signals reflected this MU. We used a blind source-separation method based on convolutive-ICA to identify a MU in the iEMG. Briefly, like ICA, this method searches for statistically independent sources that project to the sensor space. Furthermore, convolutive-ICA searches for sources which, whenever active, generate a stable waveform or activation pattern. For motor unit decomposition, the sources are MUs, and the activation patterns are their spike shapes, i.e., motor unit action potentials (MUAPs).

Applying convolutive-ICA decomposition to the first 10 s of iEMG during maximum abduction yielded one MU with an average firing rate of 4.9 Hz (MU1 in Figures 3a and 3d). A board-certified clinical neurophysiologist visually confirmed the detected MU. Spike-triggered averaging (STA) of the iEMG, i.e., averaging iEMG signals time-locked to the discovered firings, revealed significant MUAPs for both iEMG needles (Figure 3b, top, both p < 0.001). We then applied spike-triggered averaging (STA) to SQUID signals to test if the identified MU was observable in the simultaneous SQUID recordings. Indeed, for all 40 SQUID channels, the STA locked to firing moments of the iEMG MUA yielded statistically significant waveforms, i.e. motor unit action fields (MUAFs, Figure 3b, two bottom rows, all p < 0.001). Therefore, we concluded that the spiking activity of an invasively recorded MU was measurable using non-contact SQUID recordings.

Next, we asked whether we could decompose SQUID MMG signals to directly identify single MUs. Critically, the simultaneously recorded iEMG allowed us to validate putatively identified MUs. Thus, we reversed the approach and applied the convolutive-ICA to the 40-channel MMG recording. Convolutive-ICA detected two MUs with 52 and 17 spikes in the first 10 s of the maximum abduction condition (Figure 3, MU2 and MU3). We applied spike-triggered averaging of the SQUID signals to estimate the MUAFs of these MUs.

For both MUs, this yielded significant spike waveforms in all 40 MMG channels (Figure 3c, two bottom rows, all p < 0.001). As this analysis was performed only on the magnetic SQUID signals, it was possible that the identified waveforms reflected technical magnetic artifacts rather than muscular MU activity. The simultaneously recorded iEMG allowed us to validate putative MUs and to rule out potential artifacts. If the identified magnetic components reflected artifacts, the iEMG recordings should not show any MU spikes at the magnetically identified firing moments. If in contrast, iEMG recordings showed significant spikes at these firing moments, the identified MMG components reflected true MUs.

We found evidence for the latter scenario. The average iEMG signals time-locked to the magnetically identified firing moments indeed showed statistically significant MUAPs (Figure 3c top; MU2 p < 0.001 for both needles; MU3 p < 0.001 for needle 2 and p = .014 for needle 1) that were also confirmed by a board-certified clinical neurophysiologist. Thus, we concluded that MU activity can be directly decomposed from SQUID MMG.

We performed additional analyses to further characterize the MU activity identified in MMG. For each MU, the MUAFs of different SQUID channels differed in magnitude but seemed very similar in waveform shape (Figures 3b and 3c bottom row). Thus, we first addressed the question if MUAFs of a single MU reflected scaled versions of the same waveform. If that was the case, then the space spanned by the matrix of MUAFs would be one dimensional. Alternatively, MUAF of different channels could reflect different waveforms and thus span more than one dimension. To test this, we quantified the intrinsic dimensionality of MUAFs across MMG channels using PCA. We found a dimensionality of 2, 4 and 2 for MU1, MU2 and MU3, respectively (Figure 3e). Thus, for all MUs, we found that more than one waveform shape underlay the MUAFs recorded at different channels.

Second, we addressed the question if the MUs decomposed from iEMG and MMG reflected distinct MUs. In contrast, one of the two MUs decomposed from MMG (MU2 and MU3) could be the same motor unit as the MU1 detected from iEMG. For the latter scenario, firing moments of MUs should overlap with each other. To test this, we projected the entire 60 s interval of maximum abduction through the decomposition filters of the three detected MUs to identify further firing moments (Figure 4a). We quantified common firing moments as spikes of different MUs with less than 0.5 ms temporal offset. This yielded 6% and 28% of their common firing moments of MU2 and MU3 with MU1, respectively, which is below the 30% threshold typically applied for duplicated sources^11,14^. However, suboptimalities of the experimental setup (e.g. movement) could have led to missed spikes and thus to an underestimation of shared firing. Therefore, in a second test, we directly compared the waveforms of the identified MUs in the iEMG space (Figure 4b). If two components corresponded to the same MU, their MUAPs should not show any significant difference. Thus, for both MMG based MUs (MU2 and MU3), we trained an LDA classifier to dissociate their STA-based iEMG waveform from the iEMG waveform of MU1 (Figure 4b, materials and methods). We found that the waveforms of both MU2 and MU3 were significantly different from the waveform of MU1 (MU2/MU3: F1-scores: 0.81/ 0.83; both p < 0.001). We concluded that neither MU2 nor MU3 were the same as MU1.

Third, we investigated if the two MUs detected from the MMG decomposition corresponded to distinct motor units? Muscle activity generated a movement that affected the relative position of MMG sensors and muscles. This could lead to different position-dependent waveforms of one single MU, which in turn could result in a decomposition of one single MU into several components. The simultaneous iEMG recordings allowed us to test this, because the iEMG was not affected by any jitter in the relative position of MMG sensors and muscle. Thus, if the MUs identified in MMG reflected the same underlying MU, their iEMG MUAPs should have the same waveform. Alternatively, if they correspond to different MUs, their iEMG MUAPs should show a significant difference. Therefore, as for the previous analysis, we compared iEMG waveforms between MU2 and MU3 using an LDA classifier (Figure 4b, bottom two rows). We found that the iEMG waveforms of MU2 and MU3 were significantly different (F1-score = .83; p < 0.001). Therefore, we concluded that the MUs decomposed from MMG correspond to distinct MUs.

The above analyses show that the three MUs decomposed from iEMG and MMG corresponded to three different MUs with distinguishable activity patterns in iEMG. In a final analysis, we investigated if it was possible to also detect all these three MUs with distinguishable activity patterns in MMG. To this end, we tested if an LDA classifier could tell the MUAFs of MU1 apart from those of MU2 and MU3 (Figure 3b and 3c, bottom row, and Figure 4c). Indeed, LDA revealed significant differences between the magnetic activity patterns of these MUs (MU2/MU3: F1-scores = 0.71/0.83; both p < 0.001). Thus, all three MUs showed distinct MMG waveforms.

In sum, we observed strong similarities between SQUID MMG and iEMG recordings. We found that invasively identified MUs are detectable with MMG, and that automatic motor unit decomposition allows to identify invasively validated single MUs in SQUID MMG signals.

### OPM MMG

In a second experiment, we investigated whether MU decomposition is also possible for MMG recorded with OPM. We followed the same experimental approach as for the SQUID experiment, but this time recording MMG using 5 triaxial OPMs (15 channels total) together with simultaneous iEMG (Figures 5 and 6). Similar as for SQUIDs, OPM recordings showed an increase of signal power for higher levels of voluntary contraction (Figure 5d). However, although OPM recordings were conducted in a better magnetically shielded room as compared to SQUID recordings (BMSR-2, Materials and Methods), the power increase for OPM recordings was less pronounced and had a lower bandwidth as compared to SQUID recordings (Figure 2c). This is likely due to two reasons. First, the employed OPM sensors (QZFM-gen-3; QuSpin Inc.) had a much lower nominal bandwidth (135 Hz^34^) than the SQUID sensors (10 kHz). Second, OPM sensors were deployed as magnetometers with a sensitivity of 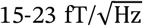, which is inferior to SQUID gradiometers with a sensitivity of 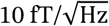.

**Figure 6.**
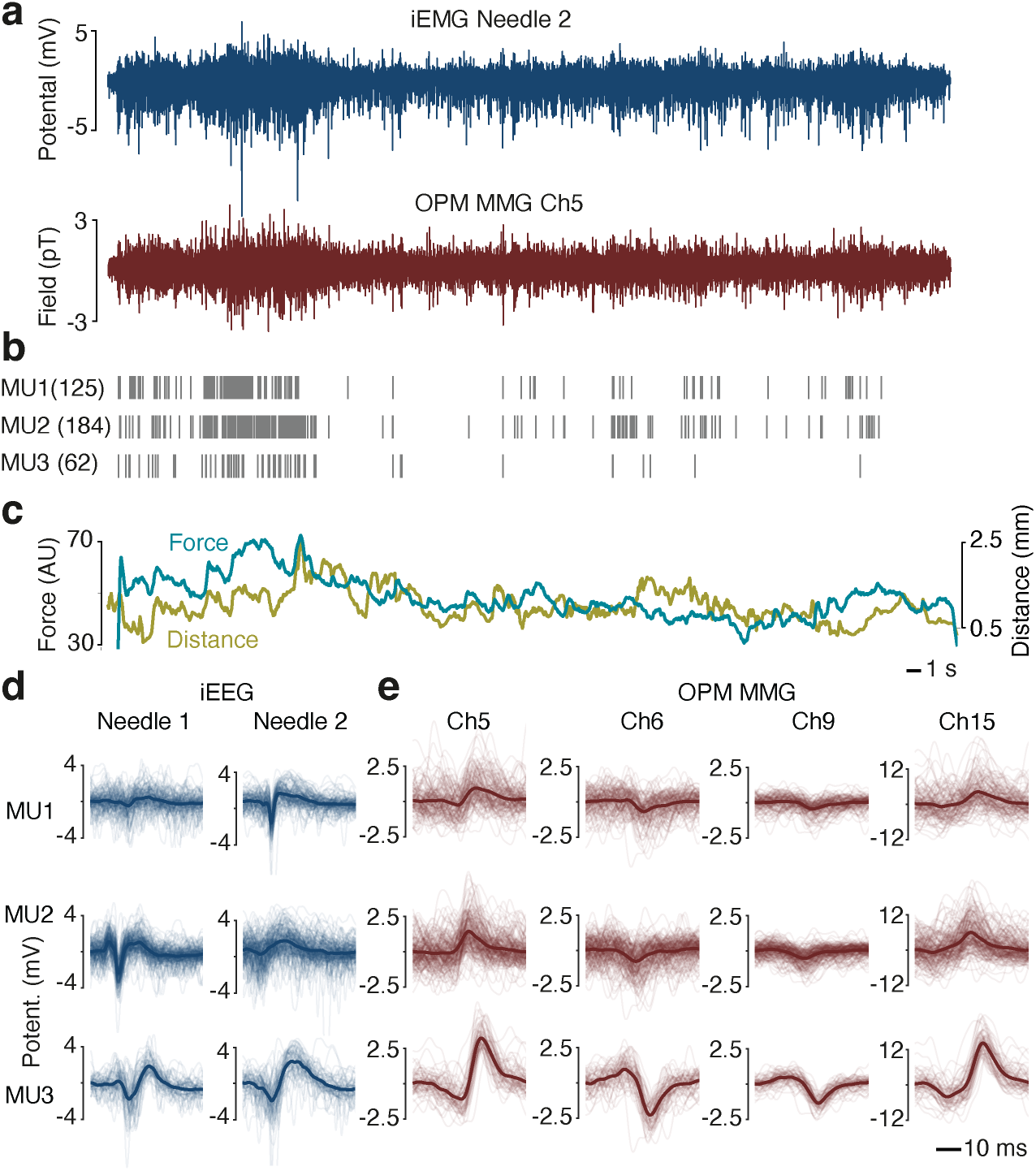
Dissociating MUs in experiment 2 **a**, iEEG (top) and OPM MMG (bottom; 1 representative channel) during the entire 60 s of maximum abduction. **b**, firing moments of the 3 identified MUs. MUs were decomposed from the first 10 s. Discharge patterns of MUs reflect the amplitude of the IEEG. **c**, Abduction force of the pinky finger and distance between the hand and the OPM positioned lateral to the hand. **d**, single discharge (thin lines) and average (thick lines) iEEG MUAPs of all identified MUs. iEEG was band-pass filtered from 20 to 1000 Hz (forward filtering). **e**, single discharge (thin lines) and average (thick lines) OPM MUAFs (4 representative channels) of all identified MUs. MMG signals are band-pass filtered from 10 to 900 Hz (forward filtering).

We next tested if, despite the lower signal fidelity, OPM MMG could be used for motor unit decomposition. First, we investigated if activities of an invasively identified single MU is observable in OPM recordings. We used convolutive-ICA to detect MUs in iEMG recordings and tested if these MUs are observable in simultaneous OPM recordings (Figure 5). Applying convolutive-ICA to the first 10 s of iEMG during maximum abduction revealed two MUs (65 and 63 spikes) with significant MUAP waveforms for both iEMG needles (MU1 and MU2 in Figure 5, p < 0.01 for both needles and MUs). We tested if these MUs were observable in the simultaneously recorded OPM MMG. Indeed, for all 15 OPM channels, the average of MMG signal locked to the iEMG firing moments showed statistically significant MUAFs (Figure 5b, two bottom rows, all p < 0.001 for both MUs). Thus, the magnetic activity of the invasively detected MUs was observable with OPM.

We next investigated if we could directly decompose the OPM MMG to identify MUs. We applied convolutive-ICA to the 15-channel OPM MMG during the first 10 s of maximum abduction. This resulted in one MU with 30 spikes (Figure 5c, MU3). For this MU, all 15 OPM channels showed significant spike waveforms (Figure 5c, right column, all p < 0.001). Critically again, we used the simultaneous iEMG to validate this component. Indeed, for both needles, the average iEMG locked to the discovered firing moments showed significant MUAPs (Figure 5c, top, both p < 0.001). Thus, the component detected in OPM MMG did not reflect magnetic artifacts, but a true MU. As for SQUID MMG, we next asked if the average waveforms of different OPM channels reflected the same waveform. Again, we applied PCA to quantify the dimensionality of the space spanned by MUAFs which resulted in 3, 2 and 3 PCs for MU1, MU2, and MU3, respectively (Figure 5e). Therefore, as for SQUIDs, we found that the spike shapes of different OPM channel reflected more than a single underlying waveform.

Furthermore, as for SQUIDs we obtained converging evidence that the three identified MUs indeed reflected distinct MUs. We identified further firings of these units in the entire 60 s recording (Figure 6b) and compared their firing moments. For all pairs, we found less than 30% overlap (MU1 with MU2/MU3: 4%/7%; MU2 with MU1/MU3: 7%/4%; MU3 with MU1/MU2: 28%/8%). Also, LDA classifiers were able to dissociate MUs from each other based on their needle activity patterns (Figure 6d; MU1 vs MU2, F1-score = 0.88, p < 0.01; MU1 vs MU3, F1-score = 0.62, p = 0.009; MU2 vs MU3, F1-score = 0.68, p < 0.001).

As for SQUIDs, we next tested if all three MU could in principle be dissociated in the OPM MMG recording. To answer this, we tested if the magnetic activity patterns of the three MUs differ from each other (Figure 6e). MU3 showed significant differences to the other two units that were identified from the iEMG decomposition (MU1 vs MU3, F1-score = 0.68, p < 0.001; MU2 vs MU3, F1-score = 0.61, p = 0.028).

However, LDA classification of the magnetic activity patterns of MU1 and MU2 that were detected from iEMG decomposition did not reach statistical significance (Figure 6e, two top rows, F1-score = 0.54, p = 0.12).

In sum, we found that, although OPM provided lower sensitivity than SQUID, invasively validated single MUs could be directly identified using OPM MMG.

The time-course of MU activity across the full 60s recording (Fig. 6b) showed a substantial variability. In a final analysis, we tested if this variability could be due to the variability of the applied abduction force and associated movements of the hand relative of OPM sensors. To test this, during the OPM experiment, we recorded the pinky finger”s abduction force and the distance between one of the OPM sensors and the subject”s ADM muscle along with OPM signals (Figure 1 and Figure 6c). Force and distance were positively correlated, indicating that abduction slightly pushed the muscle away from the OPM sensor. Moreover, we found that the low pass filtered cumulative spike train of the three discovered MUs (Figure 6b) was indeed significantly correlated with both, force and distance signals (Figure 6c, Pearson correlation coefficient: 0.57 and 0.16, for force and distance, respectively; both p < 1e-6). Thus, the variability of decomposed spike discharges could be at least partially explained by variability of the applied abduction force.

## Discussion

Here, we provide proof-of-principle evidence that single MU activity can be identified from both SQUID and OPM MMG. Our findings are based on key methodological approaches. First, we employed simultaneous invasive EMG along with MMG recordings. This provided ground truth to verify that the MUs that were non-invasively identified in MMG reflected true MUs rather than magnetic artifacts. Second, we applied an automatic decomposition method based on convolutive-ICA^14^ to detect MU activity in an unbiased and automatic fashion. Finally, we investigated our research question using two complementary approaches. On the one hand, we detected MUs in iEMG recordings and tested if these MUs were observable in SQUID and OPM recordings. On the other hand, we identified potential MUs in MMG and used the simultaneous iEMG to verify these MUs. Both approaches provided converging evidence that single MUs can be directly extracted from MMG.

In both, SQUID and OPM experiments, convolutive-ICA applied to 10 s of iEMG and MMG during maximal ADM abduction allowed us to identify one or two MUs in each modality. Firing was generally variable across time with firing rates from 4.9 to 6.5 Hz for iEMG and 1.7 to 5.2 Hz for MMG. The number of discovered MUs and their estimated firing rates were lower than in previous EMG studies^12,14,36–38^. Using a similar analytical approach, Holobar et al.^37^ decomposed surface EMG recordings of a 65 channel grid at 10% MVC (Maximum Voluntary Contraction) of ADM, and found 7 MUs with an average firing rate of 12 Hz. Farina et al.^36^ studied 8 subjects and reported that between 11 and 24 invasively detected MUs produce distinguishable patterns in a 5-by-5 bipolar EMG grid for ADM contractions below 12.5% MVC. Tanji and Kato^38^ studied 24 ADM MUs at various contraction levels with minimum and mean firing rates of 6.1 Hz and 8.3 Hz, respectively. Moreover, in most of these studies, MUs showed regular firing patterns^12,14^. The discrepancy between the present results and these previous studies is likely due to two reasons.

First, in all above-mentioned studies, EMG was recorded during isometric contractions with a constant force while hand and fingers were fixed. In contrast, in the present study the hand or fingers were not fixed, and subjects were merely asked to keep their hand position and contraction level constant without explicit position or force feedback. Consequently, hand position and force varied during the recording (Figure 6c).

On the one hand, variability of the applied force implicates a corresponding variability in the underlying MU firing patterns. Indeed, we found a significant correlation between abduction force and firing rates. On the other hand, unintentional hand movements led to a relative movement of the ADM and MMG sensors. This relative movement changes the projection pattern of MUs onto sensors. However, the employed decomposition method relies on a stable morphology of MUAPs and of their projection pattern onto sensors. Thus, the relative movement of muscle and sensors deteriorates the performance of the MMG decomposition, which decreases the number of identifiable MUs and detectable spikes of a single MU. A similar issue arises when decomposing EMG recordings from dynamic contraction paradigms, however not due to changing projections but due to changes of MUAP morphology ^39^. Indeed, as expected for stable projections of iEMG, iEMG decomposition seemed generally more stable than MMG decomposition. The decomposition patterns found for first 10 s of the data yielded higher numbers of discharges at later time points for iEMG than for MMG (MU1 vs MU2/MU3 in Figure 4a and MU1/MU2 vs. MU3 in Figure 6a).

A second reason for the difference between the present findings and previous studies is likely the different employed sensors. Specifically, many EMG studies employ dense arrays of surface EMG electrodes^14,37^, that are particularly well suited to decompose the distinct projection patterns and morphologies of different MUs. In contrast, although the present MMG recordings were performed with multiple SQUID or OPM sensors, the distance between individual sensors was much larger than for typical EEG grids, which deteriorates the ability to dissociate different MUs. Furthermore, SQUID and OPM sensors were more distant to the muscle than sEMG or iEMG electrodes, which reduces resolvable details and MU discriminability^9^. Furthermore, for the case of OPM, the low sensor bandwidth of QuSpin sensors (135 Hz) further deteriorates the situation. This likely explains why for the case of OPM, MUAF patterns of two distinct invasively recorded MUs were not distinguishable from each other.

In sum, variations of force and sensor position, the large distance of sensors between each other and to the muscle source as well as the low sensor bandwidth of OPM likely reduced the number of MUs and spikes that could be automatically extracted from MMG. However, despite these technical limitations, as a proof-of-principle the present results show that even under comparatively suboptimal and dynamic conditions non-contact MMG allows to directly extract single MU activity. On the one hand, this underscores the potential of MMG as a novel, non-contact modality to measure and investigate muscle activity. On the other hand, this points towards fruitful directions for future research and technological developments.

First, future studies may employ spatially stabilized recordings and constant forces with biofeedback to delineate the practical limits of MMG MU decomposition under optimal conditions. Second, to increase the yield of identifiable MUs, it may be particularly fruitful to increase the number and density of MMG sensors effectively moving towards HD-MMG recordings^30^. OPM sensors, which are comparatively small and can be flexibly placed are particularly promising for this approach. Furthermore, this may allow to fully exploit the unique ability of OPM sensors to record up to 3-dimensional vectorial magnetic information^30^. Third, the limited bandwidth of currently available OPM sensors provides an opportunity for further technological improvements^40–42^ that may directly benefit the application of these sensors for MMG. Forth, optimizing the parameters of the convolutive-ICA method or using alternative automatic and manual decomposition methods could potentially improve MU detection in MMG. Finally, although for the present study testing individual subjects and muscles was sufficient to provide the intended proof of principle, future studies are required to assess the variability of MU detection across the population and different muscles.

In summary, our results provide proof-of-principle evidence that invasively validated MU activity can be directly extracted from non-contact MMG recorded with SQUID and OPM sensors. Our findings underscore the potential of MMG as a non-contact modality to measure and study muscle activity in health and disease with high fidelity.

## Acknowledgements

This study was supported by ERC-AdG 2021 #101055186 and by the German Aerospace Center (DLR) with funds provided by the Federal Ministry for Economic Affairs and Climate Action (BMWK) under the grant number DLR 50WM2168.

## Author contributions

NN: Conceptualization, Methodology, Investigation, Data curation, Software, Formal analysis, Visualization, Writing – original draft, Writing – Review & Editing

JM: Conceptualization, Methodology, Investigation, Writing – Review & Editing

SH: Methodology, Investigation, Writing – Review & Editing

TM: Methodology, Funding acquisition, Investigation, Writing – Review & Editing

PB: Methodology, Investigation, Writing – Review & Editing

MS: Conceptualization, Supervision, Resources, Project administration, Funding acquisition, Writing – original draft, Writing – Review & Editing

